# Hybrid endosomal coats containing different classes of sorting nexins

**DOI:** 10.1101/2025.07.29.667382

**Authors:** Navin Gopaldass, Sudeshna Roy Chowdhury, Ana Catarina Alves, Lydie Michaillat Mayer, Véronique Comte-Miserez, Andreas Mayerv

## Abstract

Endosomes are protein sorting stations where multiple coats form tubulovesicular carriers exporting proteins to the Golgi, the plasma membrane, or endo-lysosomal compartments. Distinct classes of sorting nexins are assumed to form distinct homogeneous coats that define the endosomal sorting routes and their cargos. Snx3 and the SNX-BARs Vps5-Vps17 belong to different sorting nexin classes. They form homogeneous Retromer-dependent coats that differ in structure and their modes of membrane association and cargo recognition. Here, we show the formation of hybrid coats between purified SNX-BARs, Snx3 and their cargos. Hybrid coats assemble at variable subunit ratios and diameters and show greater membrane scaffolding activity than homogeneous coats. In vivo, Snx3 and SNX-BARs colocalise and mutually impact the sorting of their respective cargos. Although simultaneous binding of Snx3- and SNX-BARs to Retromer is sterically prohibited, hybrid coats incorporate both SNXs in a common complex, probably linked by Retromer oligomerisation. We hence propose that SNX-BARs and Snx3 form Retromer-mediated hybrid coats in novel, stoichiometrically adaptable configurations that allow to adjust endosomal carriers for transporting varying ratios of cargo.

## Introduction

Retromer is a key player in endosomal recycling. This protein complex was discovered in yeast as a pentamer, consisting of two subcomplexes: a heterodimer of the SNX-BAR sorting nexins Vps5 and Vps17, and a heterotrimer of Vps26, Vps29 and Vps35 (Seaman *et al*, 1998a). Today, the term Retromer is used to refer to this heterotrimer alone.

Retromer is recruited to endosomal membranes through SNX proteins. In yeast, Retromer binds the SNX-BAR dimer Vps5-Vps17 or Snx3 (Seaman *et al*, 1998b; Horazdovsky *et al*, 1997; Harrison *et al*, 2014; Strochlic *et al*, 2007). In mammalian cells, Retromer binds SNX3 (Harterink *et al*, 2011; Vardarajan *et al*, 2012; Chen *et al*, 2013), but it interacts with the homologues of the yeast Vps5/Vps17 dimer only indirectly. These homologues, the SNX-BARs SNX1/SNX2 and SNX5/SNX6, form the ESCPE complex (Simonetti *et al*, 2017, 2019; Lopez-Robles *et al*, 2023). Unlike the yeast SNX-BARs, ESCPE-1 interacts with Retromer through an additional sorting nexin, SNX27 (Lopez-Robles *et al*, 2023; Chandra *et al*, 2025, 2022; Guo *et al*, 2024; Simonetti *et al*, 2022).

CryoET structures of Retromer bound to either SNX-BARs or Snx3 have been solved (Leneva *et al*, 2021; Kovtun *et al*, 2018; Chen *et al*, 2025; Kendall *et al*, 2022, 2020). These structures revealed that Retromer can oligomerize to form arch-like structures that cross-link SNX proteins. They show Retromer associating with SNX3 and SNX-BARs in different ways and cross-linking them in different patterns (Figure 1C). For example, while Vps26 contacts the membrane in Snx3/Retromer coats, this is not the case in the SNX-BAR/Retromer coats, where Vps26 has no contact to lipids and binds the membrane only indirectly through the SNX-BAR layer. Both the SNX-BAR and the SNX3-based Retromer coats show a SNX layer that covers the membrane extensively but with limited regularity (Leneva *et al*, 2021; Kovtun *et al*, 2018). The binding sites for Snx3 and the SNX-BARs are positioned such that Retromer could only bind one or the other type of sorting nexin, but not both at the same time.

**Figure1:**
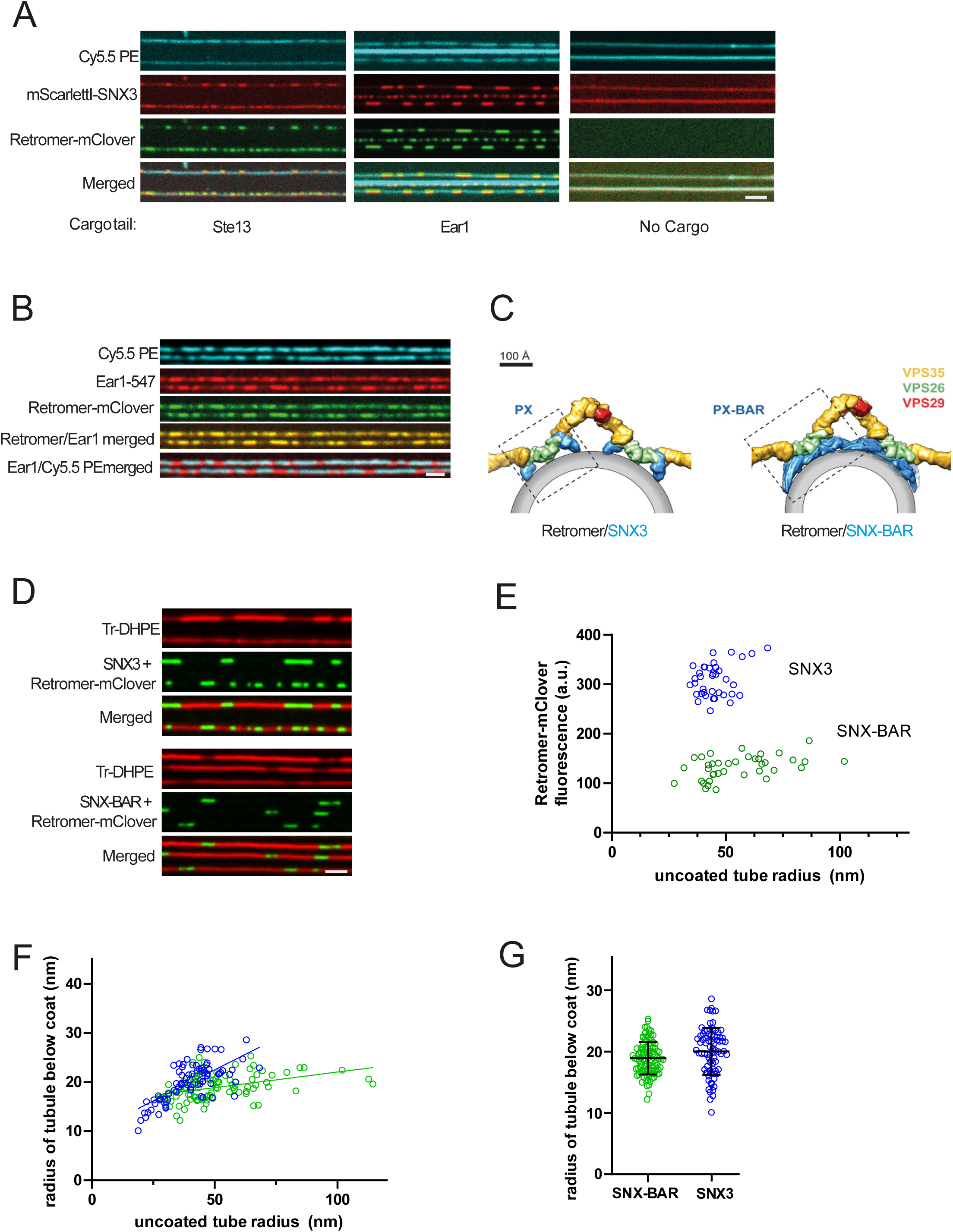
Formation of Snx3-Retromer coats and comparison with SNX-BAR-Retromer coats. **A.** Snx3-Retromer coat formation. SMTubes were formed in the presence of the lipid marker Cy5.5-PE and incubated with 10 µM cargo peptide for 10 min. Then, 50 nM of ^mScraletI^Snx3, Retromer^mClover^ (carrying mClover on Vps29) and 10 µM unlabelled cargo peptide were added. Tubes were imaged by confocal microscopy after 5 min of incubation. **B**. Cargo concentration by Snx3-Retromer coats. Tubes were formed as in A, preincubated with fluorescently labelled cargo peptide Ear1-547 for 5 min and further incubated with 50 nM Snx3, 50 nM Retromer^mClover^ and 10 µM of Ear1-547 until coat formation (2-3 min). After a brief wash with phosphate-buffered saline (PBS) buffer, tubes were imaged by spinning disc confocal microscopy. **C**. Differing modes of Retromer membrane interaction in the Snx3 and SNX-BAR coats, taken from (Leneva *et al*, 2021). **D**. Comparison of Retromer coats based on Snx3- and SNX-BARs. SMTubes, which were labelled through TexasRed-DHPE (TR-DHPE), were incubated for 5 min with 25 nM SNX-BAR/Retromer^mClover^ or 50 nM Snx3/Retromer^mClover^ and imaged as in B. **E**. Quantification of the density of Retromer-mClover on Snx3/Retromer or SNX-BAR/Retromer coats as a function of the uncoated tube radius. The uncoated tube radius was quantified through the fluorescence intensity of TR-DHPE. The fluorescence had been calibrated by comparison to SMTs constricted by Dynamin I, which constricts lipid tubules to a defined radius of 11.2 nm (Roux *et al*, 2010) . **F**. The lipid tube radius under Snx3/Retromer or SNX-BAR/Retromer coats was quantified the same approach as in E. **G**. Pooled data from F, not considering the uncoated tube radii. All scale bars are 2 µm.

In current models SNXs usually define distinct recycling pathways and contribute to the selection of specific sets of cargo proteins (Buser & Spang, 2023; Weeratunga *et al*, 2020; Weering *et al*, 2010). For example, yeast Vps10 is mainly ascribed to the Retromer/Vps5-Vps17 complex while Ear1, Ste13, Kex2 or Ftr1 are addressed by the Retromer/Snx3 complex (Strochlic *et al*, 2008, 2007; Voos & Stevens, 1998; Harrison *et al*, 2014; Suzuki *et al*, 2019). Similarly, in mammals, SNX27- and SNX3-Retromer complexes recycle specific cargo to the plasma membrane and the TGN, respectively (Temkin *et al*, 2011; Lauffer *et al*, 2010; Yang *et al*, 2018; Ghai *et al*, 2011; Lee *et al*, 2016; Steinberg *et al*, 2013; Harterink *et al*, 2011; Wassmer *et al*, 2009; Cullen & Korswagen, 2012; McGough *et al*, 2014, 2018; Tian *et al*, 2021; Cui *et al*, 2018). Sorting occurs at the endosome surface, where coat components can be enriched in domains, which generate tubulo-vesicular cargo carriers (Varandas *et al*, 2016; Thompson *et al*, 2007; Derivery *et al*, 2012, 2009; Antón-Plágaro *et al*, 2024; Puthenveedu *et al*, 2010). However, there is a priori no reason why different coat complexes might not be recruited simultaneously into the nascent endosomal carriers, since many of them are quite compatible with the highly curved surface of these tubulo-vesicular structures. This might lead to the formation of “hybrid coats”, where a single tubule contains a series of homogeneous subdomains with different SNX/Retromer coats, or to heterogeneous coats, in which different types of coat components interact and are co-recruited in the same coated domain (Gopaldass *et al*, 2024).

The concept of hybrid carriers is compatible with the fact that the spectra of cargos that depend on SNX3 or the SNX-BARs overlap, both in yeast and in mammalian cells (Bean *et al*, 2017; Steinberg *et al*, 2013). This might reflect the existence of distinct cargo exit routes that are redundant, perhaps due to similarities in the sorting signals for different SNXs (Strochlic *et al*, 2007; Ma & Burd, 2019), but it is also consistent with multiple classes of cargos and SNXs jointly using a single endosomal carrier for exiting the endosome. In mammalian cells, evidence for and against a role of common carriers incorporating SNX-BAR and SNX3 cargo has been reported. β2AR, which depends on SNX27 for returning to the plasma membrane, and Wntless, which requires SNX3 for retrograde transport to the Golgi, colocalize on the same endosomes and partition into the same leaving carriers, though with differing enrichment factors (Varandas *et al*, 2016). On the other hand, knockdown of the Drosophila orthologues of SNX1/SNX2 and SNX5/SNX6, with which SNX27 interacts and cooperates in sorting cargo (Simonetti *et al*, 2022; Guo *et al*, 2024; Simonetti *et al*, 2023; Chandra *et al*, 2025, 2022), was reported not to influence Wntless function (Harterink *et al*, 2011), with the caveat that the efficiency of the knockdown was only 80% at the RNA level and the effects on the protein levels were not tested. The same study found that mammalian Wntless enters only around 20% of SNX-BAR tubules but instead leaves endosomes in smaller SNX3-containing structures.

We set out to probe the potential for generating hybrid coats utilizing an in vitro system (Gopaldass *et al*, 2023) and the sorting nexins Vps5-Vps17 and Snx3, because structures of their respective coats have been solved (Leneva & Kovtun, 2024; Kovtun *et al*, 2018). The in vitro system offers the advantage to monitor coat growth from purified coat components on synthetic supported membrane tubes (SMTubes) in real time in an optically well-defined setting. This allows quantitative assessments of coat radius, speed of growth, and of the density of membrane coverage. Here, we exploit this system to show that yeast SNX-BARs and Snx3 readily form adaptable hybrid coats that can incorporate cargos for both classes of sorting nexins and have properties that are distinct from those of homogeneous SNX-BAR or Snx3 coats. We confirm the relevance of these findings *in vivo* by showing Retromer-mediated interaction between SNX-BARs and Snx3 and by analysing the interdependency of Snx3 and the SNX-BARs for sorting their respective cargos.

## Results

We previously described the formation of SNX-BAR and Retromer-SNX-BAR coats from purified proteins and synthetic supported membrane tubes on the surface of glass slides (SMTubes) (Gopaldass *et al*, 2023). To extend the system and ultimately enable the analysis of potential hybrid coats, we determined the conditions for Snx3 coat formation on SMTubes using recombinant Snx3 purified from E. coli. and Retromer complex (Vps26-Vps29-Vps35) and SNX-BAR (Vps5-Vps17) purified from yeast (Supplementary Figure 1) (Purushothaman *et al*, 2017; Gopaldass *et al*, 2023). As cargo is necessary to stabilize the interaction between Snx3 and Retromer (Leneva *et al*, 2021; Lucas *et al*, 2016), we synthesized small peptides corresponding to the cytosolic tails of Ste13 and Ear1, two bona fide Snx3 cargos. The peptides carried an N-terminal HIS_6_ tag to recruit them to the membrane via Ni-NTA lipids, and a C-terminal cysteine for maleimide-mediated coupling to fluorescent dyes. The coat proteins were labelled by expressing them as fusion proteins with red or green fluorescing tags, with the tags being fused to Vps17 in the SNX-BAR complex and to Vps29 in Retromer.

### Membrane scaffolding activity and Retromer saturation of Snx3 and SNX-BAR coats

While ^mScarletI^Snx3 bound to the SMTubes in the absence of the Ear1 and Ste13 cargo peptides, it required these peptides to recruit Retromer^mClover^ and form concentrated domains on the tubules (Figure 1A). The intensity of the fluorescent lipid marker Cy5.5.-PE, which had been integrated into the membrane tubules, was reduced underneath these concentrated domains. This indicates that the membrane tubes were constricted in these areas relative to the rest of the membrane, where Snx3 and Retromer did not concentrate into a denser arrangement. We hence refer to these constricted regions with a strong concentration of SNX and Retromer as coated domains. We tested whether these Snx3-Retromer domains recruited cargo using a fluorescent Alexa-547-labelled version of the Ear1 peptide (Ear1^547^) (Figure 1B). Coated domains marked by Retromer^mClover^ strongly accumulated Ear1^547^, confirming that these domains were active for cargo recruitment. All subsequent experiments with Snx3, unless stated otherwise, were performed in the presence of 10 µM of the Ear1 cargo peptide.

SNX-BARs can form dimers and higher-order associations, which at elevated SNX-BAR concentrations can suffice to scaffold a membrane tubule with these proteins alone (Gopaldass *et al*, 2023; Lopez-Robles *et al*, 2023; Weering *et al*, 2012; Sun *et al*, 2020; Zhang *et al*, 2021). By contrast, Snx3 has only a PI3P-binding PX domain and no BAR domain, and it does not show a tendency for spontaneous oligomerization. Snx3 therefore requires Retromer to form a coat and impose curvature on the membrane (Leneva *et al*, 2021)(Figure 1C). Probably for this reason the structure of the Snx3-Retromer coat shows each Snx3 molecule associated with Retromer (Harrison *et al*, 2014; Leneva *et al*, 2021), unlike the SNX-BAR-Retromer coat, which contains also SNX-BARs that are not linked to Retromer (Kovtun *et al*, 2018). We hence compared the amount of Retromer per tubule length in SNX-BAR/Retromer and Snx3/Retromer coats as a function of the initial membrane tube radius (Figure 1D and E). While the amount of Retromer^mClover^ increased with the starting tube radius for SNX-BAR/Retromer coats (Gopaldass *et al*, 2023) there was no such tendency for the Snx3/Retromer coats, also since they entirely failed to constrict wider tubes. The density of Retromer^mClover^ in the Snx3/Retromer coats was around twice the density found in the SNX-BAR/Retromer coats (Figure 1E). We also measured the lipid fluorescence of the tubules under the SNX-BAR and the Snx3 coats (Fig. 1F). This value is directly proportional to the radius of the tubule and can be calibrated by comparing the signals to those under a dynamin oligomer, which compresses the underlying membrane to a known, invariant radius (Roux *et al*, 2010; Dar *et al*, 2015). Applying this calibration revealed that Snx3 coats were on average slightly wider than the SNX-BAR coats (Figure 1G) (20 ± 3.8 nm vs. 19 ± 2.6 nm radius to the outer membrane surface), in line with previous structural studies (Leneva *et al*, 2021). However, Snx3-Retromer coats were more sensitive to the initial radius of the non-constricted tube than the SNX-BAR-Retromer coats (Figure 1F). Whereas the radius of SNX-BAR-Retromer coats increased only little with initial tube radius, the radius of the Snx3-Retromer coat increased more steeply when wider tubes had to be constricted. Thus, the Snx3-Retromer coat may be more adaptable in curvature.

Since the density of Retromer in the Snx3 coat is higher than in the SNX-BAR coat we tested whether both coats are saturated with Retromer. To this end, we incubated SNX-BAR or Snx3 with either equimolar concentrations of Retromer^mClover^ (25 nM) or with a 4- fold excess (100 nM) and measured the amount of Retromer^mClover^ incorporated into the coats for tubes of various starting radius (Figure 2 A to D). Similar amounts of Retromer^mClover^ were integrated into Snx3 coats, irrespective of whether 25 or 100 nM Retromer^mClover^ were offered (Figure 2 C and D). SNX-BAR coats integrated up to twice the amount of Retromer^mClover^ when it was present at 100 nM (Figure 2 A and B). Even at this elevated concentration, the density of integrated Retromer remained sensitive to the curvature of the membrane that must be scaffolded. It increased by another factor of two when the coat had to constrict wider membrane tubes. However, the excess of Retromer (100 nM) enabled the SNX-BARs to constrict much wider tubes than the 1:1 mixture (25 nM). This is consistent with Retromer providing driving force for coat formation (Gopaldass *et al*, 2023), because compressing wider tubes requires more work than compressing thinner tubes. Snx3-based coats behaved very differently. They could oligomerize only on much thinner tubes than the SNX-BAR coats, and even a fourfold excess of Retromer^mClover^ did not convey to them the capacity of constricting significantly wider tubes (Figure 2D).

**Figure 2:**
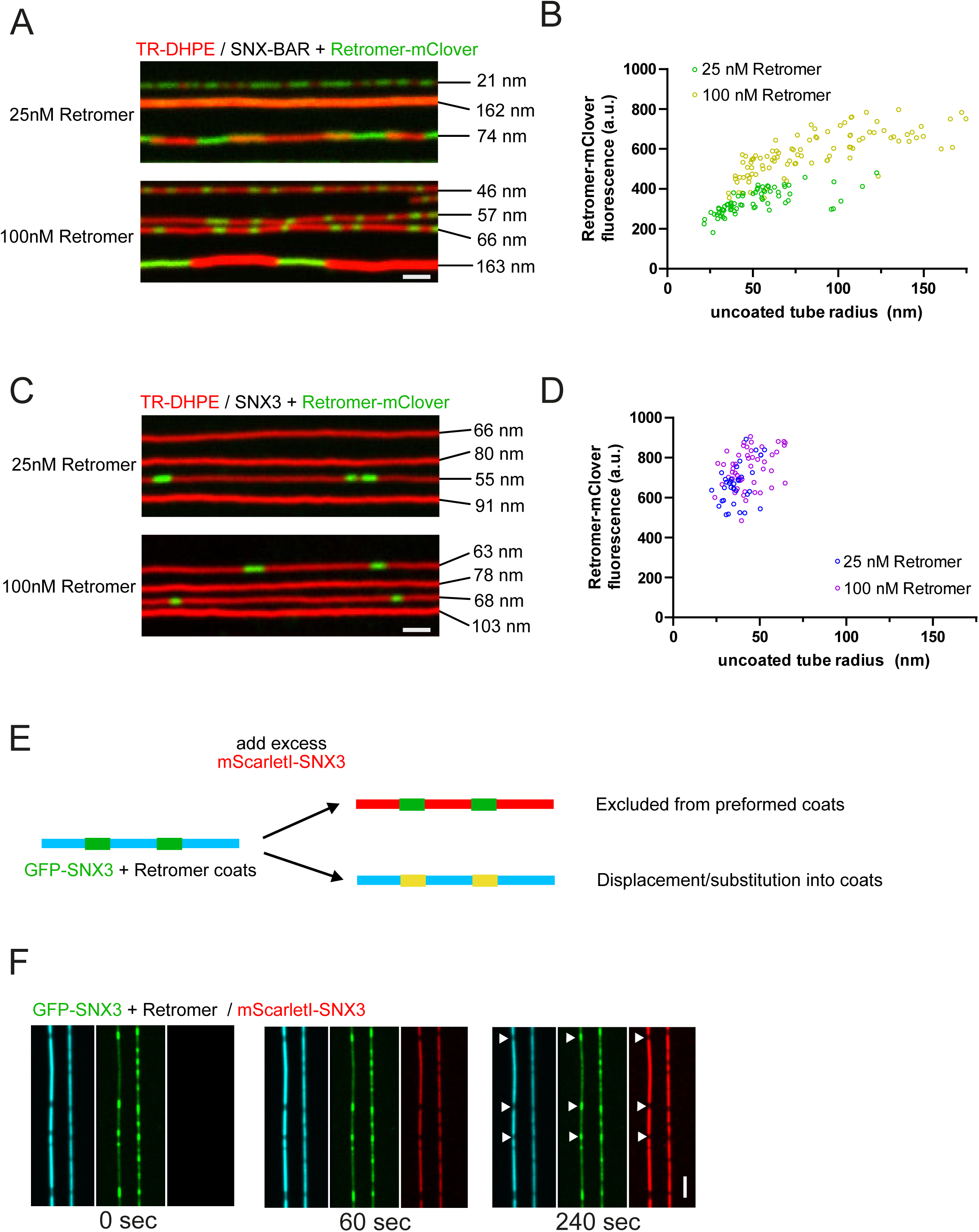
Influence of Retromer concentration on SMT constriction and coat formation by Snx3 and SNX-BARs **A.** Influence of Retromer concentration on SNX-BAR coat formation. SMTubes labelled with TexasRed-DHPE (TR-DHPE) were incubated for 5 min with 25 nM SNX-BAR and 25 nM or 100 nM Retromer^mClover^ and imaged by spinning disc fluorescence microscopy. The radius of the non-coated region of each tube is indicated. It was determined as in Fig. 1E. **B**. Quantification of Retromer^mClover^ fluorescence in coats covering SMTs of various starting radius. The radii of coated and non-coated regions were determined as in Fig. 1. Green: 25 nM SNX-BAR / 25 nM Retromer^mClover^ (n=71); yellow: 25 nM SNX-BAR / 100 nM Retromer^mClover^ (n=97). **C.** Influence of Retromer concentration on Snx3 coat formation. SMTubes were incubated for 5 min with 10 µM unlabelled ^HIS6^Ear1 cargo peptide, 25 nM Snx3 and 25 nM or 100 nM Retromer^mClover^ and imaged by confocal microscopy. The non-coated tube radius is indicated for each tube. **D**. Quantification of Retromer^mClover^ fluorescence in coats on tubes of various starting radius, determined as in B. Blue: 25 nM Snx3 / 25 nM Retromer^mClover^ (n=33), Purple: 25 nM Snx3 / 100 nM Retromer^mClover^ (n=50). **E**. Lack of subunit exchange in Snx3-Retromer coats. SMTubes were incubated for 2 min with 50 nM ^GFP^Snx3/Retromer and 10 µM ^HIS6^Ear1 cargo peptide until coats were formed. After a brief wash with buffer to remove the unbound proteins, an excess (100 nM) of ^mScarletI^Snx3 was added. Tubes were imaged by spinning disc microscopy for the indicated periods. Scale bars = 2 µm.

SNX-BAR-Retromer coats do not incorporate new SNX-BARs or exchange SNX-BARs with the pool in solution (Gopaldass *et al*, 2023). We tested this aspect also for Snx3-Retromer coats, using a two-stage experiment. In a first stage, ^GFP^Snx3-Retromer coats were formed and non-bound ^GFP^Snx3 was washed out. In a second stage, an excess of ^mScarlet-I^Snx3 was flushed into the chamber. This second wave of red fluorescing Snx3 bound readily to the non-coated regions of the tubes but was not incorporated into regions where ^GFP^Snx3 had pre-formed coats (Figure 2 E, F). Altogether, our data suggests that both SNX-BAR and Snx3-based Retromer coats are stable and do not readily exchange subunits. Nevertheless, Snx3-Retromer coats can perform less work for membrane scaffolding. They carry a fixed Snx3/Retromer ratio and are saturated with Retromer, unlike the SNX-BAR/Retromer coats, which are more adaptable in terms of their coverage by Retromer.

### SNX-BARs and Snx3 can co-integrate into a hybrid Retromer coat

Since Retromer can interact with both SNX-BARs and Snx3, we tested whether Retromer could form mixed coats incorporating both classes of sorting nexins. When SMTubes were incubated with equimolar amounts of ^mScarletI^Snx3, SNX-BAR^GFP^ and Retromer, we observed the formation of hybrid coats harbouring both ^mScarletI^Snx3 and SNX-BAR^GFP^ (Figure 3A and B). These hybrid coats could constrict tubules of up to 125 nm initial radius (Fig. 3C), resembling in this respect the SNX-BAR coats (Fig. 1F). However, the hybrid coats were larger, with a radius of 22 ± 4.7 nm (Fig. 3 D), compared to 19 ± 2.6 nm for homogeneous SNX-BAR-Retromer and 20 ± 3.8 nm for homogeneous Snx3-Retromer coats (Fig. 1G). Hybrid coat formation depended on Retromer. Co-incubating SMTubes with SNX-BAR^GFP^ and ^mScarletI^Snx3 without Retromer led to the formation of coats containing only SNX-BARs but not enriching Snx3 in this zone (Figure 3 E, F). This suggests that Snx3 is not recruited into hybrid coats by a direct interaction with SNX-BARs but requires Retromer for this. Using fluorescently labelled Ear1 and Vps10 peptides revealed that the Retromer hybrid coats concentrate both peptides (Figure 3 G, H) and are hence competent for cargo integration.

**Figure 3:**
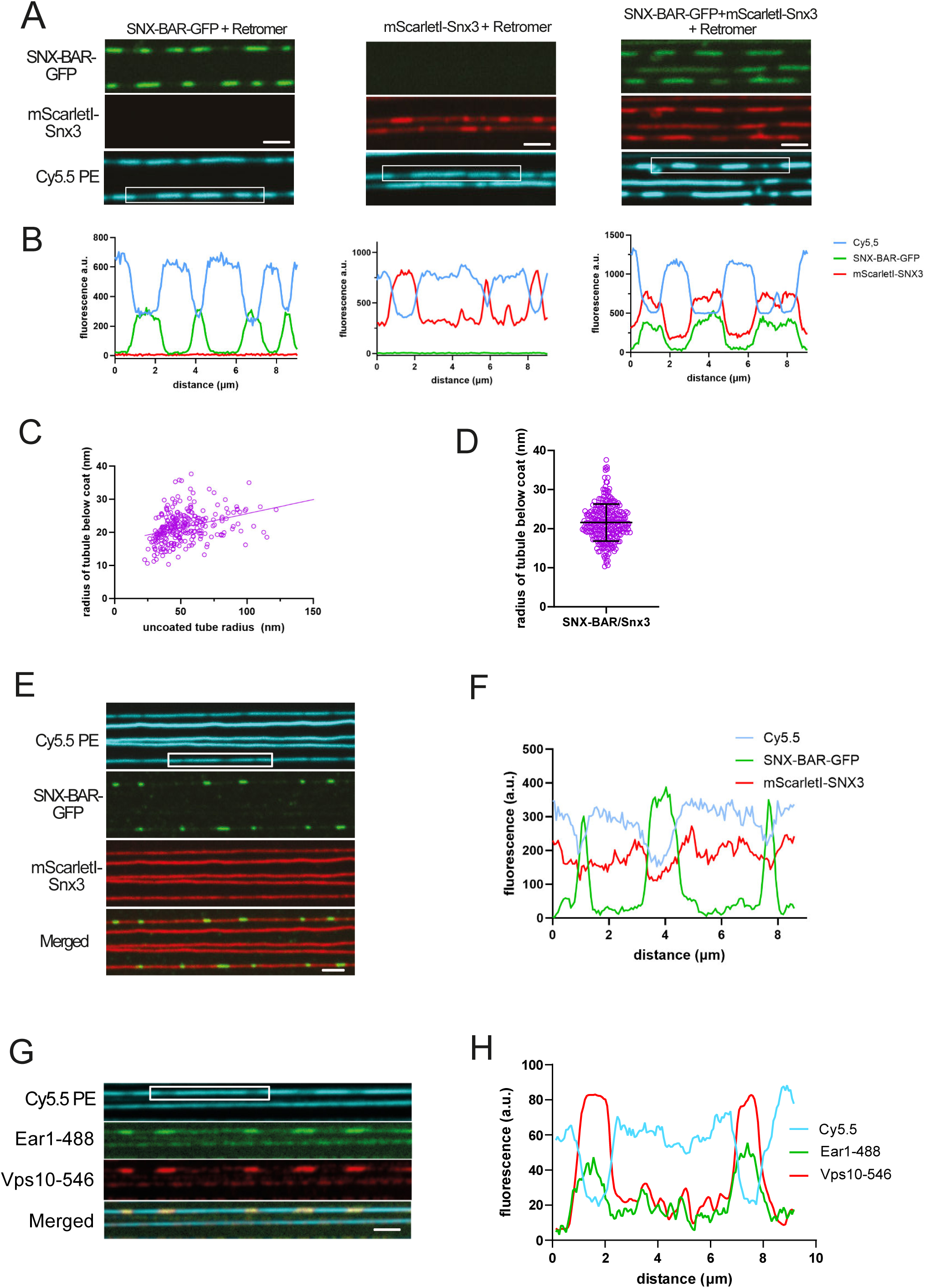
Characterization of hybrid Snx3/SNX-BAR/Retromer coats. **A**. Co-integration of Snx3 and SNX-BARs. SMTubes were pre-incubated with 10 µM ^His6^Ear1 peptide and then incubated with either 25 nM SNX-BAR^GFP^/Retromer, 50 nM ^mScarletI^Snx3/Retromer or 25 nM ^mScarletI^Snx3/SNX-BAR^GFP^/Retromer (all conditions in the presence of 10 µM ^HIS6^Ear1) for 3 to 5 min and imaged by spinning disc microscopy. **B**. Intensity scans along the tubes boxed in A. **C**. Quantification of the radius under hybrid coats as a function of the starting tube radius (n = 275). **D**. Cumulative plot of the data in C, disregarding the uncoated tube radius. **E**. Lack of hybrid coat formation in the absence of Retromer. SMTubes were incubated with 100 nM of SNX-BAR^GFP^, 50 nM of ^mScarletI^Snx3 and 10 µM ^His6^Ear1 peptide as in A, but without Retromer. **F**. Intensity scan along the tube sections boxed in E. **G**. Co-integration of Snx3 and SNX-BAR cargo. SMTubes were pre-incubated with 5 µM ^HIS6^Ear1-488 and 5 µM ^His6^Vps10-546. After 10 min of incubation with the cargo peptides, 25 nM each of Snx3, SNX-BAR and Retromer were added together with 5 µM of the labelled cargo peptides. After 3 to 5 min of incubation, unbound proteins were washed away, and tubes were imaged by confocal microscopy. **H**. Intensity scan along the boxed tube section shown in C. All scale bars: 2 µm.

The structures of Snx3 or SNX-BAR coated membrane tubules show limited regularity of these coats and some voids, which might provide space to incorporate other proteins (Leneva *et al*, 2021; Kovtun *et al*, 2018). We hence performed staged experiments to test whether Snx3 could integrate into preformed SNX-BAR-Retromer coats or vice versa (Figure 4 A to D). SMTs were incubated with 25 nM SNX-BAR^GFP^ and Retromer to allow the formation of coats that were 1-2 µm long. Unbound proteins were washed away and an excess (100 nM) of ^mScarletI^Snx3 was added (Figure 4A and B). After 5 min of incubation, ^mScarletI^Snx3 was mainly localized to the uncoated regions, but it did not integrate into the preformed SNX-BAR-Retromer coats. Similarly, pre-formed ^mScarletI^Snx3-Retromer coats could not integrate an excess of SNX-BAR^GFP^ (Figure 4C and D). These observations suggest that the pre-formed homogeneous coats do not contain suitable voids and binding sites and/or are not flexible enough to integrate the other sorting nexin class.

**Figure 4:**
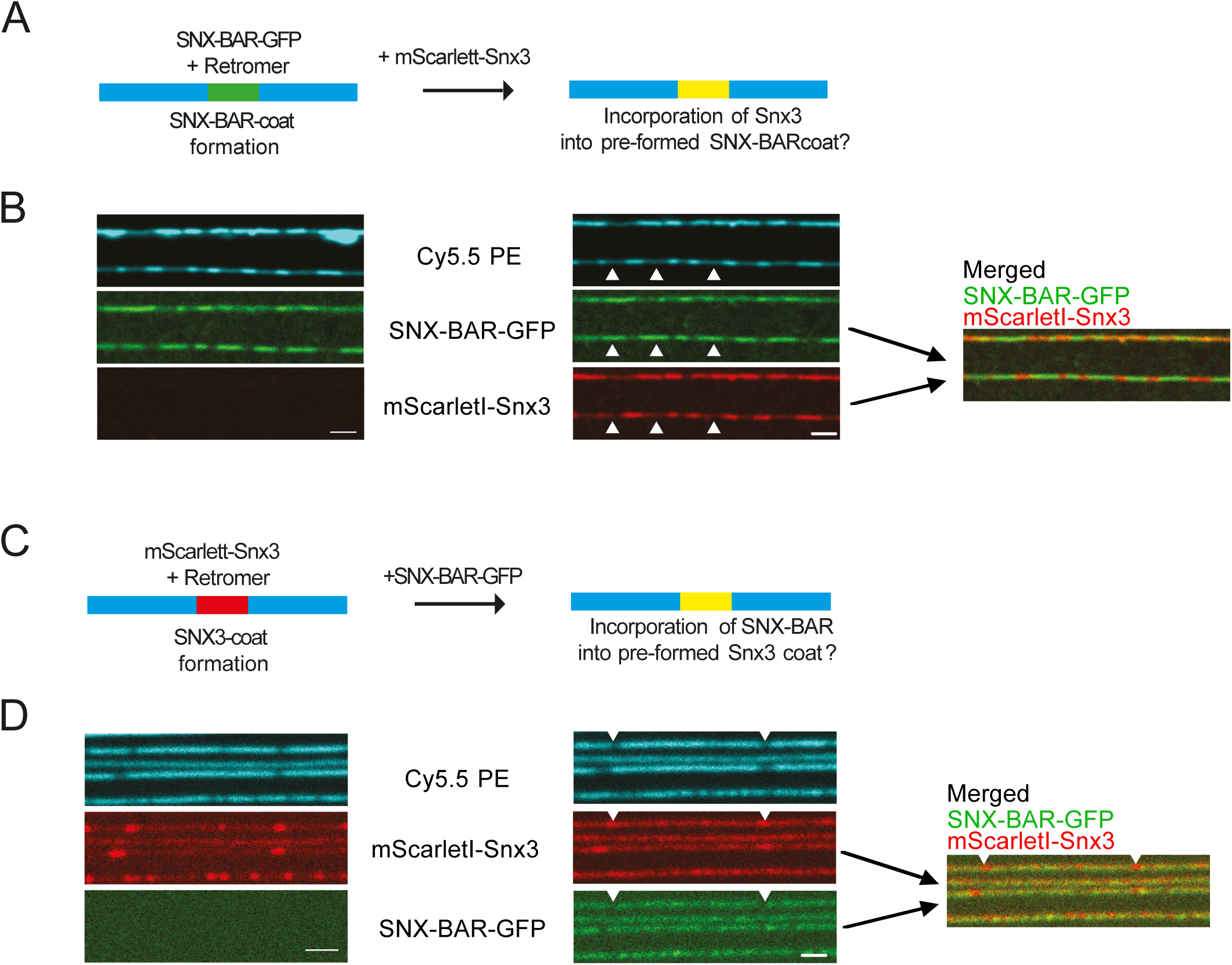
Homogeneous Snx3 or SNX-BAR coats cannot integrate the other sorting nexin class. **A.** Non-integration of Snx3 into pre-formed SNX-BAR Retromer coats. SMTubes were incubated with 25 nM of SNX-BAR^GFP^/Retromer for 2-3 minutes until coats were formed. After a brief wash to remove unbound proteins, 100 nM ^mScarletI^Snx3 was added and incubated for 5 min. Tubes and coats were imaged by confocal microscopy. Arrowheads point to SNX-BAR^GFP^/Retromer coats, which are not accessible to ^mScarletI^Snx3. **B.** Non-integration of SNX-BARs into pre-formed Snx3 Retromer coats. SMTubes were incubated with 50 nM of ^mScarletI^Snx3/Retromer for 2-3 minutes until coats were formed. After a brief wash with buffer, 50 nM SNX-BAR^GFP^ was added and incubated for 5 min. Tubes and coats were imaged by confocal microscopy. Arrowheads point to ^mScarletI^Snx3/Retromer coats, which are not accessible to the excess of added SNX-BAR^GFP^. Scale bars = 2 µm.

To directly compare SNX densities in the hybrid coats, we created a ^GFP^Snx3 construct using the same GFP tag as in our SNX-BAR^GFP^. Retromer coats were generated with either ^GFP^Snx3 or SNX-BAR^GFP^ in the presence or absence of non-tagged versions of the respective other sorting nexin (Figure 5 A, B). The overall sorting nexin concentration was kept constant (50 nM for pure coats and 25 nM of each sorting nexin class for hybrid coats, both in the presence of 50 nM Retromer) (Figure 5 A to C). The mean GFP fluorescence per tube length in the ^GFP^Snx3/Retromer coat (404 ± 80 a.u., n=40) was about twice the mean GFP fluorescence in the SNX-BAR^GFP^ coat (230 ± 40 a.u., n=40) (Figure 5 C). Considering that the SNX-BAR complex is a heterodimer of a labelled Vps17^GFP^ and an unlabelled Vps5 it follows that the SNX densities in Snx3-based and SNX-BAR-based Retromer coats are similar.

**Figure 5:**
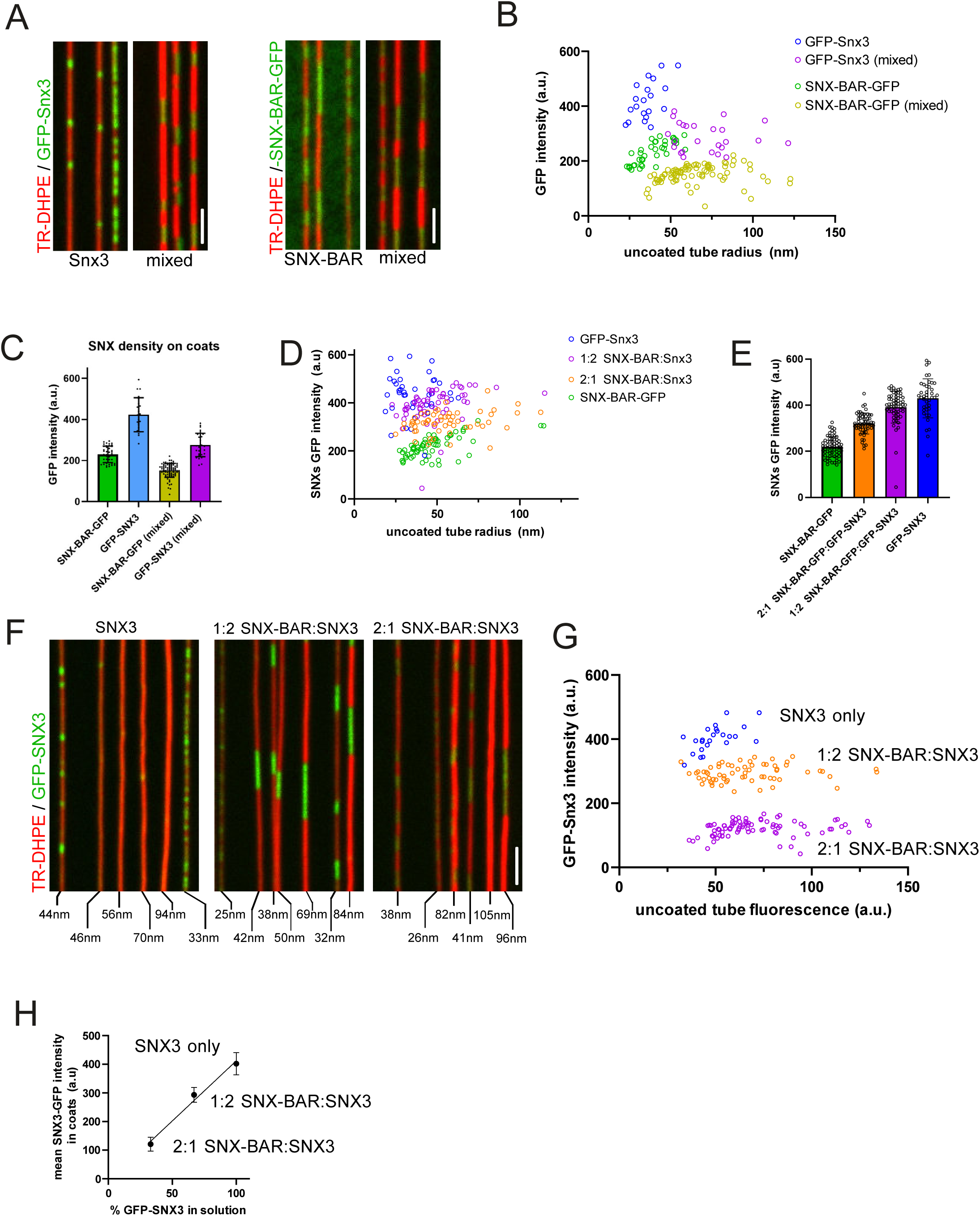
Sorting nexin stoichiometry in hybrid coats. **A**. Comparison of Snx3 and SNX-BAR densities in hybrid and the respective homogeneous coats. SMTubes were incubated with Retromer and the following combinations of sorting nexins carrying a GFP-tag either on Snx3 or the SNX-BARs: 50 nM ^GFP^Snx3 and 50 nM Retromer, 25 nM of the SNX-BAR^GFP^ heterodimer (25 nM Vps5 and 25 nM Vps17^GFP^) and 25 nM Retromer, 25 nM ^GFP^Snx3 with 25 nM SNX-BARs and 50 nM Retromer, or 25 nM Snx3 with 25 nM SNX-BAR^GFP^ and 50 nM Retromer. After 2 to 5 min of incubation, the SMTubes were imaged by spinning disc microscopy. **B**. Quantification (performed as in Fig. 1) of the GFP intensity for coats on tubes of various starting radius, comparing homogeneous SNX coats with coats formed from mixtures of the sorting nexins. ^GFP^Snx3/Retromer (n = 22), SNX-BAR^GFP^/Retromer (n = 40), ^GFP^Snx3/SNX-BAR/Retromer (n = 25), Snx3/SNX-BAR^GFP^/Retromer (n = 84) **C**. The mean ^GFP^SNX intensity for each type of coat was calculated from the data in A and B. **D**. Integration of sorting nexins at variable stoichiometries. SMTubes were incubated with 50 nM Retromer and sorting nexins at three different ratios of ^GFP^Snx3 to SNX-BARs: 1:0 = 25 nM ^GFP^Snx3 only; 1:2 = 12.5 nM SNX-BAR and 25 nM ^GFP^Snx3; and 2:1 = 50 nM SNX-BAR and 25 nM Snx3. After coat formation, unbound protein was washed away with buffer and tubes were imaged as in A. The non-constricted radius is indicated for each tube. **E**. Quantification of D. ^GFP^Snx3 in coats formed in the presence of various ratios of competing SNX-BARs was quantified as in Fig. 1. Snx3 (n = 24), 1:2 SNX-BAR:Snx3 (n = 65), 2:1 SNX-BAR:Snx3 (n = 83). **F**. Mean ^GFP^Snx3 intensity of the data in E, shown as a function of % of total SNXs in solution. Scale bars: 2 µm.

The different degrees of labelling of ^GFP^Snx3 (every subunit is labelled) and SNX-BAR^GFP^ (a heterodimer, in which only Vps17^GFP^ is labelled, but not Vps5) could also be exploited in replacement experiments. To this end, Retromer coats were generated at different ratios of ^GFP^Snx3 over SNX-BAR^GFP^ (Figs. 5 D, E). The total GFP signal in the coats should remain constant if one ^GFP^Snx3 molecule replaced a SNX-BAR dimer, but it should increase if two Snx3 molecules replaced a SNX-BAR dimer. Regarding a coat region of 6 sorting nexins as an example, this unit would contain 3 GFP when made of only of SNX-BAR^GFP^ heterodimers. In a coat assembled at a SNX-BAR^GFP^ : ^GFP^Snx3 ratio of 2:1, this unit would contain 4 GFPs (2 from Vps17^GFP^, 2 from ^GFP^Snx3), at a 1:2 ratio it would contain 5, and in a pure ^GFP^Snx3 coat it would contain 6. The fluorescence signals of hybrid coats that assembled from ^GFP^Snx3 and SNX-BAR^GFP^ at these ratios followed exactly this expected pattern (Figure 5D and E). This result suggests that one Snx3 subunit substitutes one SNX-BAR subunit and that hybrid coats can assemble at variable SNX-BAR/Snx3 stoichiometries. This latter point was further supported when we compared coats formed from mixtures containing ^GFP^Snx3 and unlabelled SNX-BARs at ratios of either 2:0, 2:1 or 1:2. Retromer was kept at the same concentration in all cases (Figure 5 F - H). The Snx3^GFP^ signal in the coat decreased in proportion to the SNX-BAR:Snx3^GFP^ ratio in solution. Hybrid coats thus do not have a fixed composition, suggesting that they are adaptable as a function of available Snx3 and SNX-BARs and/or their cargos.

### Hybrid coat formation enhances the capacity for membrane scaffolding

We noticed that hybrid coats could form on and constrict tubes of higher starting radius than homogeneous Snx3 coats (Fig. 5B, D, G), suggesting that mixed coats are more potent in deforming the membrane and that SNX-BARs might help drive Snx3/Retromer coat formation. This points to a potential synergy between Snx3 and the SNX-BARs in scaffolding the tubes. We used giant unilamellar vesicles (GUVs) to test whether this is the case. On GUVs, coat formation probably requires more work for membrane deformation because a tubular membrane conformation is not pre-defined as in the case of SMTubes. Tubulation also reduces the volume of the GUV and necessitates extrusion of luminal liquid, requiring additional work. When GUVs were incubated with 100 nM ^mScarletI^Snx3 and Retromer, tubules did not form (Figure 6). Substituting a quarter of the Snx3 by SNX-BARs (25 nM SNX-BAR^GFP^ : 75 nM Snx3) resulted in vigorous tubule formation on about 60% of the GUVs. This tubulation was stronger than the tubulation observed with a 25 nM SNX-BAR-GFP or with 100 nM ^mScarletI^Snx3 alone. Thus, hybrid coat formation allows Snx3 and SNX-BARs to provide higher membrane scaffolding activity than in respective homogeneous coats.

**Figure 6:**
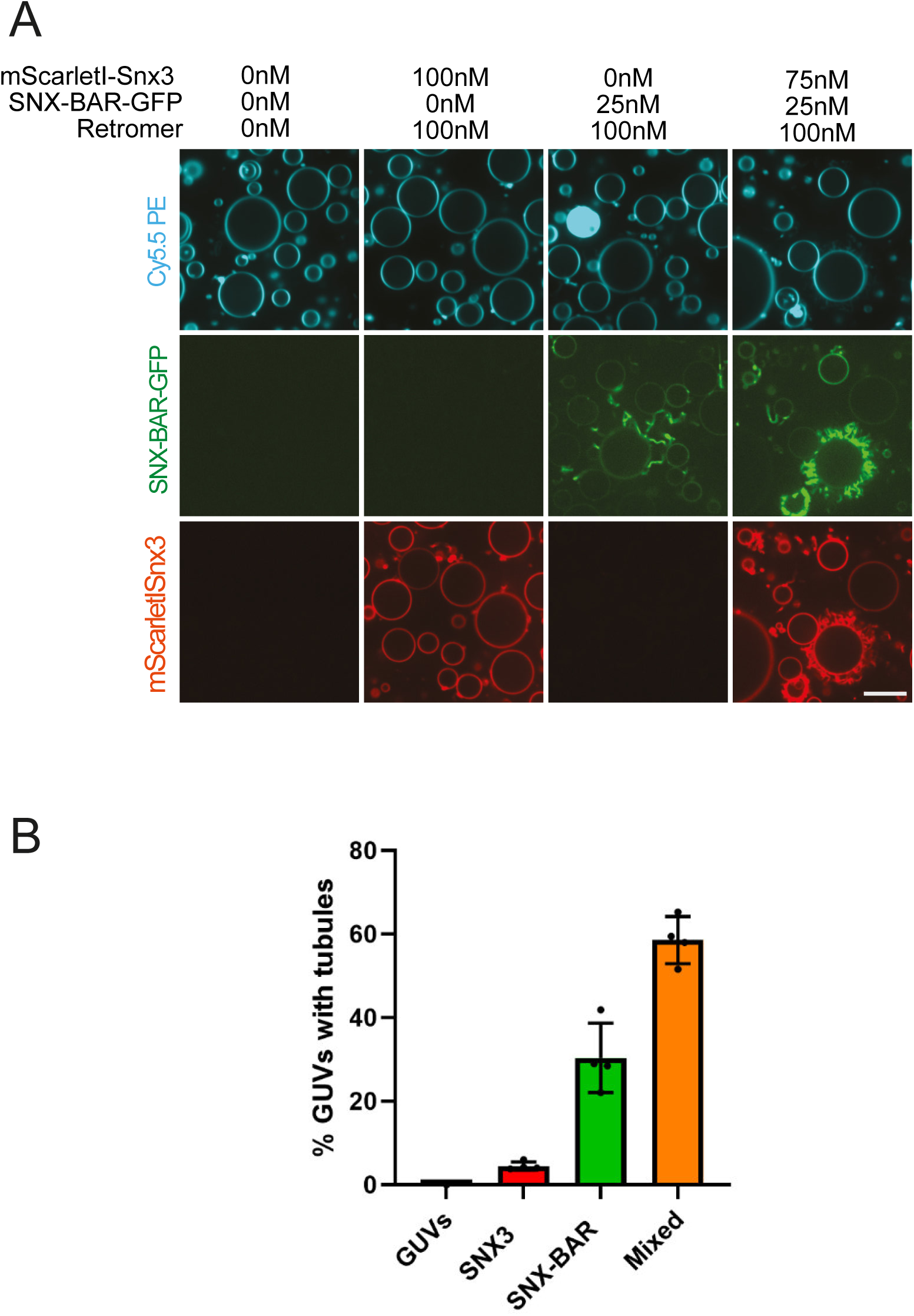
Tubulation of GUVs by hybrid coats. **A**. GUVs labelled with Cy5.5-PE were incubated with the indicated concentrations of Retromer, ^mScarletI^Snx3, and SNX-BAR^GFP^, and 10 µM Ear1 peptide. After 1 hour of incubation, the GUVs were imaged by confocal microscopy. **B**. Quantification of the percentage of GUVs with tubules for each condition shown in A. (GUVs only n=186, Snx3/Retromer n=177, SNX-BAR/Retromer n=193, Snx3/SNX-BAR/Retromer n=358). Scale bar: 5 µm.

### Hybrid coat formation in vivo

To search for in vivo correlates of hybrid coat formation, we first compared the localization of Snx3 and SNX-BARs by tagging Snx3, Vps5 or Vps17 with either mNeonGreen (mNG) or yomCherry (Figure 7A, B). The major pool of Snx3^mNG^ colocalized with Vps5^yomCherry^ (Figure 7A). To have a reference point, we performed the same experiment with Vps5^yomCherry^ and Vps17^mNG^ as these proteins form a stable heterodimer (Seaman *et al*, 1998a; Seaman & Williams, 2002) and can therefore provide a benchmark for maximal colocalization (Figure 7A). The Pearson correlation coefficients showed similar colocalization between Snx3 and Vps5^yomCherry^ as between Vps5^yomCherry^ and Vps17^mNG^ (Figure 7B).

**Figure 7:**
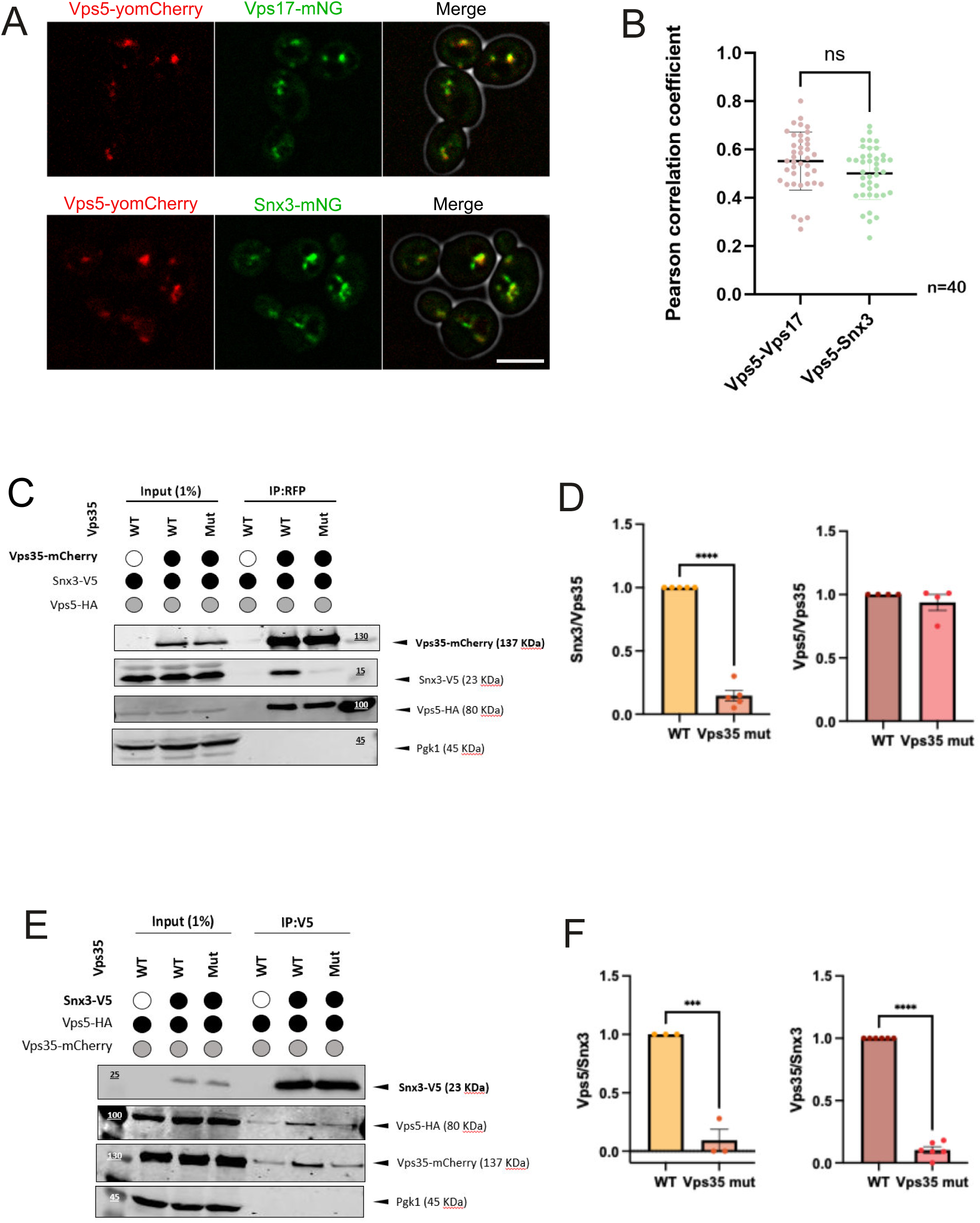
Snx3 and SNX-BARs interact in vivo depending on Retromer. **A**. Colocalization of Snx3 and SNX-BARs. Cells co-expressing Vps5^yomCherry^ and Vps17^mNG^, or Vps5^yomCherry^ and ^mNG^Snx3 were analysed by a z-series of confocal images taken at a z-distance of 0.3 µm. Average projections were generated in ImageJ. Scale bar: 5 μm. **B**. Pearson Correlation Coefficient (PCC) values were calculated from the data in A for 40 cells of each strain. Bars represent the mean values and standard deviation. **C.** Immunoprecipitation of mCherry tagged Vps35^wt^ or Vps35^QGRE^. Logarithmically growing *vps35Δ* cells expressing Vps35^mCherry^ (WT) or Vps35^QGRE-mCherry^ (Mut) from a plasmid, and genomically tagged Snx3^V5^ and Vps5^HA^, were lysed in detergent. Proteins were pulled down with an affinity matrix to mCherry and analysed by SDS-PAGE and Western blotting against the indicated tags. **D.** Quantification of the amount of Vps5 and Snx3 pulled down by mCherry-labelled Vps35^wt^ or Vps35^QGRE^. The ratios between the signals for Vps35 and Snx3 or Vps5 were calculated. For Vps35^wt^ the ratio was set to 1 as a reference. **E.** Snx3^V5^ was pulled down in an experiment as in C, using *vps35Δ* cells expressing Vps35^mCherry^ (WT) or Vps35^QGRE-mCherry^ (Mut) from a plasmid and genomically tagged Snx3^V5^ and Vps5^HA^ in the indicated combinations. **F.** Quantification of Vps35^mCherry^ and Vps5^HA^ pulled down by Snx3^V5^. The ratios between the signals for Snx3^V5^ and Vps35^mCherry^ or Vps5^HA^ were calculated. For Vps35^wt^ the ratio was set to 1 as a reference. Scale bars: 5 µm.

The binding sites of Snx3 and SNX-BARs on Retromer overlap (Leneva *et al*, 2021; Kovtun *et al*, 2018), rendering the simultaneous binding of Snx3 and SNXBARs to a single Retromer complex unlikely. However, Retromer forms arch-like dimers, which might permit binding of SNX-BARs at one end and of Snx3 at the other. To test this, we generated a strain carrying tagged SNX-BAR (Vps5^HA^), Snx3 (Snx3^V5^) and Retromer (Vps35^yomCherry^). We also generated a *vps35* mutant impaired in Snx3 binding (*vps35^QGRE^*, with the substitutions Q181A, G182A, R185A, and E186A) based on previous interaction studies (Harrison *et al*, 2014). We used immunoadsorption assays to probe whether Snx3 and SNX-BARs interact through Retromer. In the immunoadsorption experiments, wildtype Vps35^yomCherry^ pulled down both Snx3^V5^ and Vps5^HA^. By contrast, V*ps35^QGRE^*^-yomCherry^ lost the interaction with Snx3^V5^ but maintained the interaction with Vps5^HA^ (Figure 7 C, D). When we pulled down Snx3^V5^ from VPS35^yomCherry^ wildtype cells, both Vps35^yomCherry^ and Vps5^HA^ co-adsorbed to the beads (Figure 7 E, F). The Snx3^V5^-Vps5^HA^ interaction was reduced in the *vps35^QGRE^* background. This suggests that Retromer can bridge Snx3 and SNX-BARs.

To examine the in vivo relevance of this association, we tagged yeast Retromer cargo proteins and followed their fate in vivo (Figure 8). We used Vps10 as a bona fide SNX-BAR cargo (Bean *et al*, 2017; Purushothaman & Ungermann, 2018; Suzuki *et al*, 2019), and Ear1 and Ste13 as Snx3 cargos. In wildtype cells, a Vps10^mNG^ fusion protein localized to cytosolic puncta which, based on previous work (Suzuki *et al*, 2019; Day *et al*, 2018), should represent yeast endosome/late Golgi (Figure 8 A). In SNX-BAR knockout cells (*vps5Δ* or *vps17Δ*), Vps10^mNG^ colocalized with the vacuolar marker FM4-64. This vacuolar localization results from a failure to recycle Vps10 to the Golgi (Nothwehr *et al*, 1999; Cereghino *et al*, 1995). In *snx3Δ* knockout cells, Vps10^mNG^ localization was moderately affected, as evident from cells showing weak staining of the vacuolar membrane with Vps10^mNG^, in line with previous results (Bean *et al*, 2017). The Increased colocalization between Vps10^mNG^ and the vacuolar lipid marker FM4-64 confirmed this limited mislocalization to the vacuolar membrane (Fig. 8 B). Furthermore, also the *vps35^QGRE^*mutant, which targets the Retromer-Snx3 interaction, showed a partial accumulation of Vps10^mNG^ on the vacuolar membrane, suggesting that the recruitment of Snx3 to Retromer is relevant for Vps10 sorting (Supplementary Figure 2). Vice versa, the Snx3 cargos Ear1^mNG^ (Figure 8 C, D) and Ste13^mNG^ (Figure 8 E, F) showed significant mislocalization to vacuoles not only in *snx3Δ*, but also in *vps5Δ* or vps17Δ cells, suggesting that the Snx3 recycling pathway requires SNX-BARs for its function. These in vivo observations are in line with our in vitro findings and support the notion that Retromer can link SNX-BARs and Snx3 into hybrid coats that promote recovery of cargos for both classes of sorting nexins.

**Figure 8:**
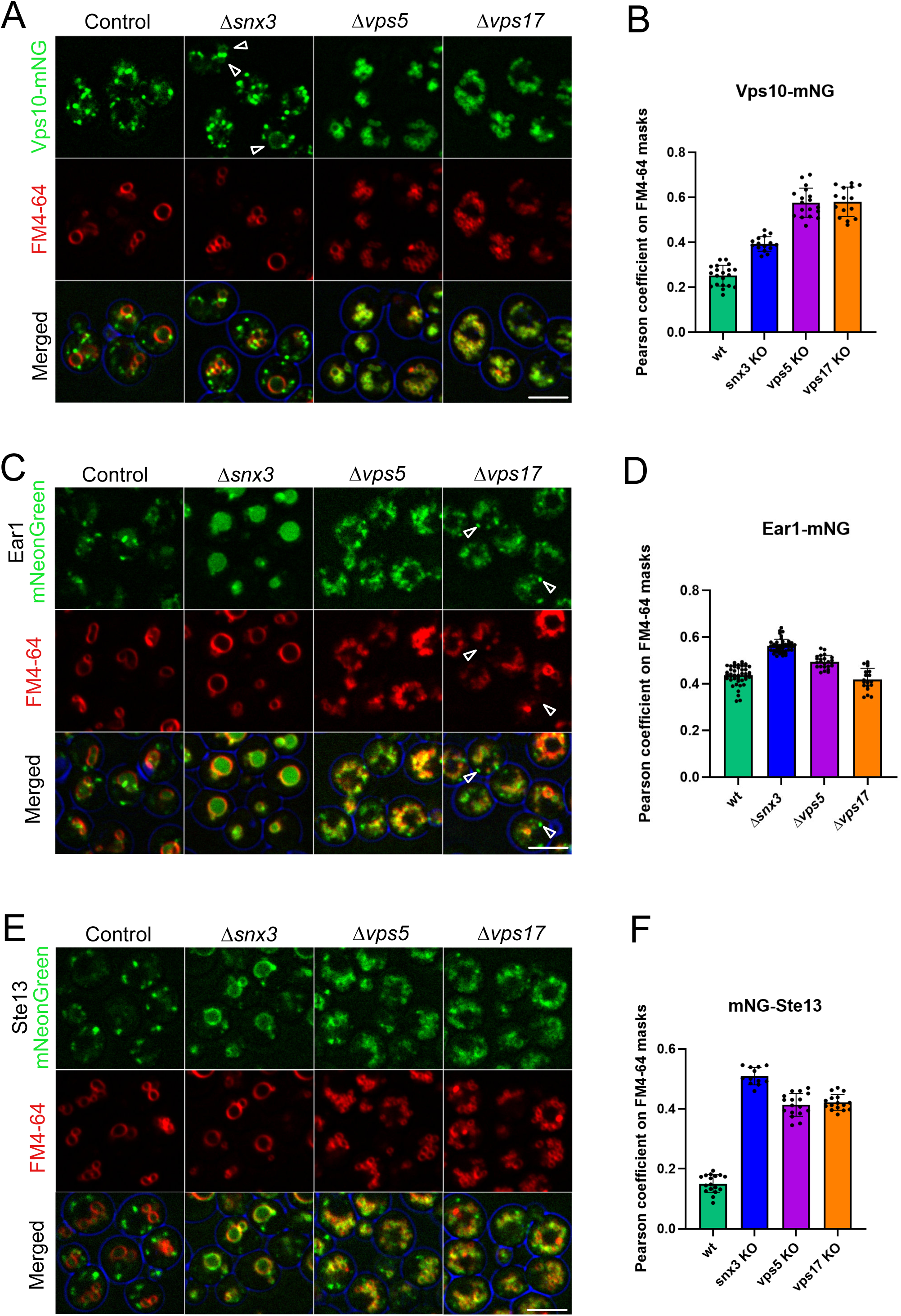
Mutual dependence of Snx3 and SNX-BARs for cargo recycling. **A**. Vps10 distribution. Logarithmically growing cells expressing Vps10^mNG^ in wildtype, *Snx3*Δ, *vps5*Δ, and *vps17*Δ background were stained with FM4-64 and analysed by confocal microscopy. Average intensity projections of confocal z-stacks taken at a z-interval of 0.3 µm are shown. **B.** Quantification of data from A. Pearson correlation coefficients values were calculated using masks defining a zone close to the FM4-64 signal around the vacuolar membrane. At least 100 cells were analysed for each strain. Data are presented as mean values +/- standard deviation. **C**. Same experiment as in A, but with cells expressing Ear1^mNeonGreen^ instead of Vps10^mNeonGreen^. Arrow heads point to Ear1 localized to endosomal structures. **D.** Quantification of data from C, performed as in B. **E**. Same experiment as in A, but with cells carrying Ste13^mNeonGreen^ instead of Vps10^mNeonGreen^. **F.** Quantification of data from E, performed as in B. Scale bars: 5 µm.

## Discussion

Structural studies revealed many features of Retromer associated with either the SNX-BAR Vps5 or Snx3 (Leneva *et al*, 2021; Kovtun *et al*, 2018; Lucas *et al*, 2016; Kendall *et al*, 2020, 2022) . These coats share the organization of Retromer dimers into arches that cross-link a SNX layer on a membrane tubule. The available structures support the current view that each SNX that associates with Retromer defines a separate sorting route. However, larger scale analyses of the impact of various endosomal coat components on the steady state distribution of a broad spectrum of cargo have revealed a significant overlap in the cargos affected by different classes of sorting nexins, including Snx3 and SNX-BARs, both in yeast and in mammalian cells (Bean *et al*, 2017; McNally *et al*, 2017; Steinberg *et al*, 2013). Further observations exist that are consistent with the formation of hybrid coats between Snx3 and SNX-BARs but were not explicitly interpreted from this perspective. In mammalian cells, Retromer cargo for SNX3 (Wntless) and the SNX-BAR-associated protein SNX27 (β2-andrenergic receptor) (Harterink *et al*, 2011; Simonetti *et al*, 2022) were partially found in and left from the same endosomal sorting domains although they use different sorting nexins and have different destinations (Varandas *et al*, 2016). The sorting signals of Wntless and β2-andrenergic receptor could be exchanged without impairing their entry into endosomal sorting domains (Varandas *et al*, 2016). Also in yeast, Snx3 and Vps17 were reported to partially co-localise, and both were necessary for efficient sorting of the iron transporter Ftr1 (Strochlic *et al*, 2007). Another link is provided by overexpression of the Snx3 cargo Ear1, which led to missorting of the SNX-BAR cargo Vps10 to the vacuole, and by the observation that Ear1 retrieval was compromised in a *vps5Δ* mutant (Suzuki *et al*, 2019). These studies were not designed to exclude indirect effects and were hence not interpreted in the context of hybrid coat formation. Therefore, models linking one type of homogenous sorting nexin coat with a defined population of cargo and a transport direction have prevailed in the general discussion of endosomal protein sorting. Our direct observation of the hybrid coat formation from purified proteins now provides a physical correlate and a potential explanation for the overlaps in the coat dependencies of cargo transport.

The hybrid coats between Snx3 and SNX-BARs that we have analysed can integrate cargo peptides for both sorting nexin classes, suggesting that they are functional for cargo collection. However, they have properties that are distinct from those of the respective homogeneous coats. While both Snx3 and SNX-BARs oligomerize with Retromer into stable coats on SMTubes, Snx3/Retromer cannot constrict wider tubules, which SNX-BAR-Retromer readily scaffolds. This difference might result from the different capacity to self-interact. SNX-BARs self-interact and form a lattice on their own that can deform the membrane, and Retromer adds additional bonds and driving force to this process (Gopaldass *et al*, 2023; Simunovic *et al*, 2015; Lopez-Robles *et al*, 2023). In a Snx3 coat, however, the sorting nexins do not self-interact and Retromer is the only element that crosslinks them and imposes curvature on the membrane (Leneva *et al*, 2021; Kovtun *et al*, 2018). Integration into hybrid coats apparently cures this lack of scaffolding activity of Snx3, allowing it to populate and constrict tubes of similar width as a homogeneous SNX-BAR coat, and to tubulate GUVs that it would otherwise not be able to deform.

Available structures of Snx3- and SNX-BAR Retromer coats suggest that simultaneous binding of the two sorting nexin classes to Retromer might be impossible due to steric clashes (Leneva *et al*, 2021; Kovtun *et al*, 2018; Lucas *et al*, 2016). Hybrid coats could thus form in two ways, which are not mutually exclusive. They might contain interspersed homogeneous zones that are covered only by Snx3-Retromer or SNX-BAR-Retromer and are too small to be resolved by our light-microscopic analysis. Alternatively, Retromer oligomers, which can form through dimerization of Vps35 or Vps26 (Lucas *et al*, 2016; Leneva *et al*, 2021; Kovtun *et al*, 2018; Kendall *et al*, 2022, 2020), could crosslink Snx3 and SNX-BARs into a joint, oligomeric structure. This latter possibility is favoured by the Retromer-dependent interaction of Snx3 with the SNX-BARs that we have detected.

The sorting nexin densities of homogeneous Snx3- and SNX-BAR-based coats on SMTs are comparable. After their formation, these homogeneous coats do not readily exchange subunits. They cannot integrate sorting nexins from the respective other class, suggesting that they are stable structures and that they do not contain suitable voids to convert into hybrid coats. The composition of the coats is apparently defined when they are being formed. During formation, however, there is considerable adaptability. Hybrid coats can form at variable Snx3/SNX-BAR ratios. Our replacement experiments suggest that this leaves the overall sorting nexin density unchanged, implying that SNX-BAR and Snx3 subunits occupy similar average membrane areas in the coats. The adaptability of hybrid coat composition is an interesting feature, particularly in combination with a role for cargo in coat formation. Snx3, for example, can bind to membranes by itself, but its integration into coats requires cargo (Fig. 1)(Lucas *et al*, 2016; Leneva *et al*, 2021). An adaptable coat stoichiometry combined with cargo dependence could offer a simple mechanism for adjusting coat composition to the amounts of cargos for Snx3 and SNX-BAR that must be transported.

The formation of hybrid coats comes with challenges for the specificity of sorting and targeting to distinct final locations. When cargos with different destinations enter the same carrier, sorting will have to occur after the formation of the carriers. Several possibilities have been proposed (Varandas *et al*, 2016). They may include correction through subsequent trafficking steps that would remove the mis-delivered cargo, or selective retention at the correct location, e.g. by interactions with resident proteins or special biophysical properties, such as the length of a transmembrane domain that must be adapted to the membrane thickness of the target compartment (Sharpe *et al*, 2010; Watson & Pessin, 2001; Cosson *et al*, 2013). These additional challenges may be offset by benefits. As we have shown above, for example, the hybrid coats can enhance scaffolding activity of Snx3, enabling it to form coats on less curved membranes, which do not permit this for a homogeneous Snx3-Retromer coat, presumably because it cannot provide the required driving force. We consider it as likely that further combinations of hybrid coats exist. This may apply to Retromer, because, in yeast, Snx3 and Snx4 localisations are distinct, but both proteins colocalize substantially with the SNX-BAR Vps17 (Strochlic *et al*, 2007). On a broader scale, it appears conceivable that also other endosomal coat components might co-enrich on tubules that are generated on the surface of a endosomes. Many of them are based on sorting nexins that have affinity for the highly curved environment of a tubular membrane (Teasdale & Collins, 2012; Weering *et al*, 2010). Thus, as soon as tubules are generated, e.g. powered by the activity of motors or actin polymerisation, they might attract a mix of coat components. Testing this possibility will require careful live imaging analyses in vivo and in vitro and continued structural analyses of the coat complexes.

## Material and methods

### Materials

Lipids were purchased from Avanti Polar Lipids (USA): Egg L-alpha-phosphatidylcholine (EPC); 1,2-dioleoyl-sn-glycero-3-phospho-L-serine sodium salt (DOPS); 1,2-dioleoyl-sn-glycero-3-phospho-(1’-myo-inositol-3’-phosphate) (PI3P). All lipids were dissolved in chloroform and stored under argon at -20°C and used within 2 months. Phosphatidylinositol phosphates were dissolved in chloroform/methanol/water (20:9:1). Texas red DHPE (Thermofisher cat. T1395MP) was purchased as a mixed isomer. The para isomer was separated by thin layer chromatography as previously described (Dar *et al*, 2015).

### Cell culture, strains and plasmids

BY4742 yeast cells were grown at 30°C in YPD (**% w/v yeast extract, **% peptone, 2 % dextrose) medium. Genes were deleted by replacing a complete open reading frame with a marker (Janke *et al*, 2004; Güldener *et al*, 1996) (see Appendix Tables 1-3 for a list of strains, plasmids and PCR primers used in this study). Gene tagging was performed as described (Sheff & Thorn, 2004). Strains used for expression and purification of the Retromer complex have been described previously (Gopaldass *et al*, 2024).

### Live microscopy

Vacuoles were stained with FM4-64 essentially as described (Desfougères *et al*, 2016) . An overnight preculture in HC medium was used to inoculate a 10 ml culture. Cells were then grown in HC at 30 °C and 150 rpm to an OD_600_ between 0.6 and 1.0. The culture was diluted to an OD_600_ of 0.4, and FM4-64 was added to a final concentration of 10 µM from a 10 mM stock in DMSO. Cells were labeled for 60 min with FM4-64, washed three times in fresh media, and then incubated for 60 min in media without FM4-64. Just before imaging, cells were concentrated by a brief low-speed centrifugation and placed on a glass microscopy slide overlaid with a 0.17 mm glass coverslip. Z-stacks were taken with a spacing of 0.3 µm and assembled into maximum projections. Imaging was performed with a NIKON Ti2 spinning disc confocal microscope with a 100x 1.49 NA lens and two Photometrics Prime BSI cameras. Image analysis was performed with ImageJ as described in (Gopaldass *et al*, 2023) for SMTubes or a python-based app. This app is available under the following link:*********. Pearson’s correlation coefficient was used to quantify the colocalization between Vps10 and FM4-64. The Nikon NIS-Elements Software Pearson’s correlation tool was used on at least 5 stacks containing at least 100 cells each. All performed experiments were repeated at least three times. SEM calculation and plotting were done with PRISM Graphpad software.

### Protein immunoprecipitation and western blotting

The interaction between SNX3, Retromer, and VPS5 was examined using a co-immunoprecipitation (co-IP) protocol adapted from (Suzuki *et al*, 2019). Yeast strains endogenously expressing Snx3-V5, Vps35-yomCherry, and Vps5-HA were inoculated in fresh YPD from saturated pre-cultures, grown over-night at 30°C and harvested at mid-logarithmic phase (OD₆₀₀ = 0.8–1.0). A total of 50 mL of culture was collected and washed twice with immunoprecipitation (IP) buffer (20 mM HEPES-KOH, pH 7.2, 0.2 M sorbitol, 50 mM potassium acetate, 1× protease inhibitor cocktail (200 µM pefablock, 11 µM leupeptin, 500 µM o-phenanthroline, 7 µM pepstatin A), 1 mM PMSF, followed by centrifugation to pellet the cells.

Cell lysis was performed using 0.5-mm zirconia beads in 300 µL IP buffer at 4°C for 5 minutes at 2500 rpm on an IKA^®^ VIBRAX^®^ VXR orbital shaker. Subsequently, 300 µL of IP buffer containing either 1.0% Triton X-100 (for Vps35 immunoprecipitation) or 1.5% Triton X-100 (for Snx3 immunoprecipitation) was added to reach final detergent concentrations of 0.5% and 0.75%. Lysates were incubated on a rotating wheel at 4°C for 2 hours, followed by clarification through centrifugation at 500 × g for 5 minutes and then at 17,500 × g for 10 minutes. Protein concentrations in the supernatants were quantified using a NanoDrop spectrophotometer and the absorption at 280 and 205 nm. 10 mg of total protein was used for each immunoprecipitation reaction. Samples were incubated with pre-equilibrated anti-RFP magnetic particles (RFP-Trap Magnetic Particles, Chromotek) or anti-V5 magnetic agarose beads (V5-Trap Magnetic Agarose, Chromotek) for 2 hours at 4°C. After incubation, beads were washed three times with IP buffer containing the appropriate Triton X-100 concentration (0.5% or 0.75%). Prior to the final wash (performed with detergent-free IP buffer), the beads were transferred to fresh microcentrifuge tubes. Bound proteins were eluted by boiling the beads at 95°C for 10 minutes in 4× NuPAGE sample buffer supplemented with 10 mM DTT, 2 mM PMSF, and 1× PIC. Protein samples were subsequently resolved by SDS-PAGE on freshly cast 4–18% polyacrylamide gels.

### Protein purification

TAP-tagged Retromer complex was extracted from yeast as previously described (Purushothaman *et al*, 2017; Purushothaman & Ungermann, 2018). Briefly, a 50 mL preculture of cells was grown for 24 hours in YPGal medium. The next day, two 1L cultures in YPGal were inoculated with 15 mL of preculture and grown for 20h at 30°C and 150 rpm to late log phase (OD_600_ = 2 to 3). All following steps were performed at 4°C. Cells were pelleted and washed with 1 pellet volume of cold RP buffer (Retromer Purification buffer: 50 mM Tris pH 8.0, 300 mM NaCl, 1 mM MgCl_2_, 1 mM PMSF, and home-made protein inhibitor cocktail (200 µM pefablock, 11 µM leupeptin, 500 µM o-phenanthroline, 7 µM pepstatin A). Pellets were either processed immediately or flash-frozen in liquid nitrogen and stored at -80°C. For cell lysis, the pellet was resuspended in one volume of RP buffer and passed once through a French press (One shot cell disruptor, Constant Systems LTD, Daventry, UK) at 2.2 Kpsi. DNase I was added to the lysate (final concentration 0.1 mg/mL) followed by a 20 min incubation on a rotating wheel. The lysate was precleared by centrifugation for 30 min at 45’000 x g in a Beckman JLA 25.50 rotor and cleared by a 60 min centrifugation at 150’000 x g in a Beckman Ti 60 rotor. The cleared supernatant was passed through a 0.2 µm filter and transferred to a 50 mL Falcon tube. 1 mL IgG beads suspension (GE Healthcare, cat 17-0969-01) was added to the supernatant. After 60 min incubation on a rotating wheel, beads were spun down and washed 3 times with RP buffer and resuspended in 2 mL RP buffer. 250 µg of home purified HIS-TEV protease from E. coli was added to the beads. After 30 min incubation at 4°C, beads were centrifuged, the supernatant containing purified Retromer subcomplex was collected and concentrated on a 100 kDa cutoff column (Pierce™ Protein Concentrator PES, 100K MWCO). The concentrated protein fraction was re-diluted in RP buffer and reconcentrated 3 times. This final step allowed for removal of TEV protease and a high enrichment of intact complexes. Proteins were concentrated to ∼2 mg/mL, aliquoted in 10 µL fractions and flash-frozen in liquid nitrogen. Proteins were stored at -80°C and used within 3 months. Thawed aliquots were used only once.

### Peptide synthesis and labelling

Peptides were synthesized at the Protein and Peptide Chemistry Facility, University of Lausanne (Switzerland) and analysed by mass spectrometry. The homogeneity of all peptides was >90%, as indicated by analytical HPLC. For fluorescent labelling, 3 to 4 mg of peptides were dissolved in 100 µL DMSO and incubated at 20°C for 12 hours with an equimolar amount of maleimide coupled fluorophore (Alexa Fluor™ 488 or 546 C5 Maleimide, Invitrogen). Labelled peptides were then purified by HPLC.

**Cargo Peptide sequences**: numbering indicates the amino acid position in the full-length protein. The sequence in brackets at the end was added to permit labelling with a maleimide coupled fluorophore.

Ste13: HHHHHHGGGG-^78^MRPRRESFQFNDIENQH

Ear1: HHHHHHGGGG-^475^GKKIINEEINLDSL-(GGC)

Vps10: HHHHHHGGGGGG-^1423^GGFARFGEIRLGDDGLIE-(GGC)

### Supported membrane tubes

SMTubes were generated as previously described (Dar *et al*, 2015; Gopaldass *et al*, 2024). Briefly, glass coverslips were first washed with 3 M NaOH for 5 min and rinsed with water before a 60 min treatment with piranha solution (95% H_2_SO_4_ / 30% H_2_O_2_ 3:2 v/v). Coverslips were rinsed with water and dried on a heat block at 90°C. Coverslips were then silanized with 3-glycidyloxypropyltrimethoxysilane (Catalogue no. 440167, Sigma) for 5 h under vacuum, rinsed with acetone and dried. Polyethylene glycol coating was performed by placing the coverslips in a beaker containing PEG400 (Sigma) at 90°C for 60 h. Coverslips were washed with distilled water and stored for up to 2 months at room temperature in a closed container.

To generate supported membrane tubes, lipids were mixed from 10 mg/mL stocks in a glass vial and diluted to a final concentration of 1 mg/mL in chloroform. The same lipid mix was used throughout this study (5% PI3P, 15% DOPS, 0.1% Texas red DHPE, 79.5% egg-PC). Lipids were then spotted (typically 1 µL, corresponding to about 1 nmol) on the coverslips and dried for 30 min under vacuum. The coverslip was mounted on an IBIDI 6-channel µ-slide (µ-Slide VI 0.4. IBIDI, Cat.No: 80606). Lipids were hydrated for 15 min with buffer (PBS pH 7.2: 137 mM NaCl, 2,7mM KCl, 10 mM Na_2_HPO_4_, 1,8 mM KH_2_PO_4_)) and SMTs were generated by injecting PBS into the chamber using an Aladdin Single-Syringe Pump (World Precision Instruments, model n°. AL-1000) at a flow rate of 1,5 mL/min for 5 min. SMTs were left to stabilize without flow for 5 min before the start of the experiment. Protein stocks (typically 1-2 µM) were first diluted in PBS and then injected in the chamber at a flow rate of 80 µL per minute. Tubes were imaged with a NIKON Ti2 spinning disc confocal microscope equipped with a 100x 1.49 NA objective.

### Quantification of SMT fluorescence

SMT fluorescence was quantified with ImageJ as described previously (Gopaldass *et al*, 2023). Briefly, line scan analysis was performed along tubules using ImageJ. For each line scan, a Gaussian curve was fitted, and the maximum height was extracted. Maximum height was then plotted against the tube length for all channels. For quantification of the radius of the tubes, lipid fluorescence values of a tubule underneath a constricted protein domain, extracted from the series of line scans described above, was sorted in ascending order. The curve typically showed two plateaus, the lower corresponding to the constricted state and the higher to the non-constricted one. Plotting the corresponding FP values confirmed that the FP-labeled protein localized to the constricted zone. For each tube, the zones corresponding to the constricted and non-constricted areas were determined manually and the mean fluorescence value was used to calculate the tube radius or the FP intensity in the corresponding region.

---

**Appendix table 1.** Strains used in this study.

| Strain | Genotype | Reference |
| --- | --- | --- |
| CUY105 | *MATa his3Δ200 leu2Δ0 met15Δ0 trp1Δ63 ura3Δ0* | EUROSCARF |
| CUY100 | *MATalpha his3Δ200 leu2Δ0 lys2Δ0 met15Δ0 trp1Δ63 ura3Δ0* | EUROSCARF |
| CUY9228 | CUY105, *VPS5pr::natNT1-GAL1pr VPS17pr::kanMX-GAL1pr VPS5::TAP-URA3 vps35::HIS3* | (Purushotham an et al, 2017) |
| CUY9495 | CUY100, *VPS26pr::HIS3-GAL1pr VPS29pr::natNT2-GAL1pr VPS35pr::hphNT1-GAL1 VPS35::TAP-kanMX vps5::TRP1 vps17::LEU2* | (Purushotham an et al, 2017) |
| CUY9711 | CUY105, *VPS5pr::natNT1-GAL1pr VPS17pr::kanMX-GAL1pr VPS5::TAP-URA3 vps35::HIS3 VPS17::GFP-hphNT1* | (Purushotham an et al, 2017) |
| CUY9932 | CUY105, *VPS26pr::HIS3-GAL1pr VPS29pr::natNT2-GAL1pr VPS35pr::hphNT1-GAL1 VPS29-mClover::kan VPS26::TAP-URA3 vps5Δ::TRP1* | (Purushotham an et al, 2017) |
| CUY9935 | CUY105, *VPS26pr::HIS3-GAL1pr VPS29pr::natNT2-GAL1pr VPS35pr::hphNT1-GAL1pr VPS29-mCherry::kan VPS26::TAP-URA3 vps5Δ::TRP1* | (Purushothama n et al, 2017) |
| SEY6210 | *MATα leu2-3,112 ura3-52 his3-Δ200 trp1-Δ901 suc2-Δ9 lys2-801; GAL* | Scott Emr |
| SEY6211 | *MATa leu2-3,112 ura3-52 his3-Δ200 trp1-Δ901 ade2-101 suc2-Δ9; GAL* | Scott Emr |
|  | *SEY Ear1-mNG (URA), vps35Δ, LEU::pRS405 Vps35prom-Vps35-mCherry-ADHterm, Vps5-3HA (TRP)* | This study |
|  | *SEY Ear1-mNG (URA), Snx3-V5, Vps5-3HA (TRP)* | This study |
|  | *SEY Ear1-mNG (URA), vps35Δ, LEU::pRS405 Vps35prom-Vps35-mCherry-ADHterm, Snx3-V5, Vps5-3HA (TRP)* | This study |
|  | *SEY Ear1-mNG (URA), vps35Δ, LEU::pRS405 Vps35prom-Q181G182R185E186-Vps35-mCherry-ADHterm, Snx3-V5, Vps5-3HA (TRP)* | This study |
|  | *SEY Ear1-mNG (URA), vps35Δ* | This study |
|  | *SEY Ear1-mNG (URA), vps35Δ, LEU::pRS405 Vps35prom-Vps35-mCherry-ADHterm* | This study |
|  | *SEY Ear1-mNG (URA), vps35Δ, LEU::pRS405 Vps35prom-Q181G182R185E186-Vps35-mCherry-ADHterm* | This study |
|  | *SEY Ear1-mNG (URA), Snx3-V5* | This study |
|  | *SEY Ear1-mNG (URA), vps35Δ, LEU::pRS405 Vps35prom-Vps35-mCherry-ADHterm, Snx3-V5* | This study |
|  | *SEY Ear1-mNG (URA), vps35Δ, LEU::pRS405 Vps35prom-Q181G182R185E186-Vps35-mCherry-ADHterm, Snx3-V5* | This study |
|  | *SEY Ear1-mNG (URA), vps35Δ, LEU::pRS405 Vps35prom-Vps35-mCherry-ADHterm, Snx3-V5, vps17Δ* | This study |
|  | *SEY Ear1-mNG (URA), vps35Δ, LEU::pRS405 Vps35prom-Vps35-mCherry-ADHterm, snx3Δ* | This study |
|  | *SEY Vps10-mNG, vps35Δ, URA::pRS406 Vps35prom-Vps35-mCherry-ADHterm* | This study |
|  | *SEY Vps10-mNG, vps35Δ, URA::pRS406 Vps35prom-Vps35-Q181G182R185E186-mCherry-ADHterm* | This study |
|  | *SEY Vps10-mNG, vps35Δ, URA::pRS406 Vps35prom-Vps35-mCherry-ADHterm, snx3Δ* | This study |
| NG450 | *SEY6210 snx3Δ::kanMX4* | This study |
| AM4134 | *SEY6211 vps17Δ::natNT2* | This study |
| SRC10 | *SEY6210 Δvps5 Vam10-V5::LEU2* | This study |
| SRC37 | *SEY6210 Δvps5 Vam10-V5::LEU2 Vps10-mNeonGreen* | This study |
| SRC39 | *SEY6210 Δvps5 Vam10-V5::LEU2 Ear1-mNeonGreen* | This study |
| SRC41 | *SEY6210 Δvps5 Vam10-V5::LEU2 mNeonGreen-Ste13* | This study |
| SRC42 | *SEY6211 vps17Δ::natNT2 Vps10-mNeonGreen* | This study |
| SRC44 | *SEY6211 vps17Δ::natNT2 Ear1-mNeonGreen* | This study |
| SRC46 | *SEY6211 vps17Δ::natNT2 mNeonGreen-Ste13* | This study |
| SRC62 | *SEY6210 snx3Δ::kanMX4 Vps10-mNeonGreen* | This study |
| SRC64 | *SEY6210 snx3Δ::kanMX4 Ear1-mNeonGreen* | This study |
| SRC66 | *SEY6210 snx3Δ::kanMX4 mNeonGreen-Ste13* | This study |
| SRC30 | *SEY6210 Vps5-yomCherry ::SpHIS5 Vps17-mNeonGreen ::CaURA3* | This study |
| SRC31 | *SEY6210 Vps5-yomCherry ::SpHIS5 Snx3-mNeonGreen ::CaURA3* | This study |
| SRC22 | *SEY6210 Vps10-mNeonGreen* | This study |
| SRC23 | *SEY6210 Ear1-mNeonGreen* | This study |
| SRC26 | *SEY6210 mNeonGreen-Ste13* | This study |

**Appendix table 2:**
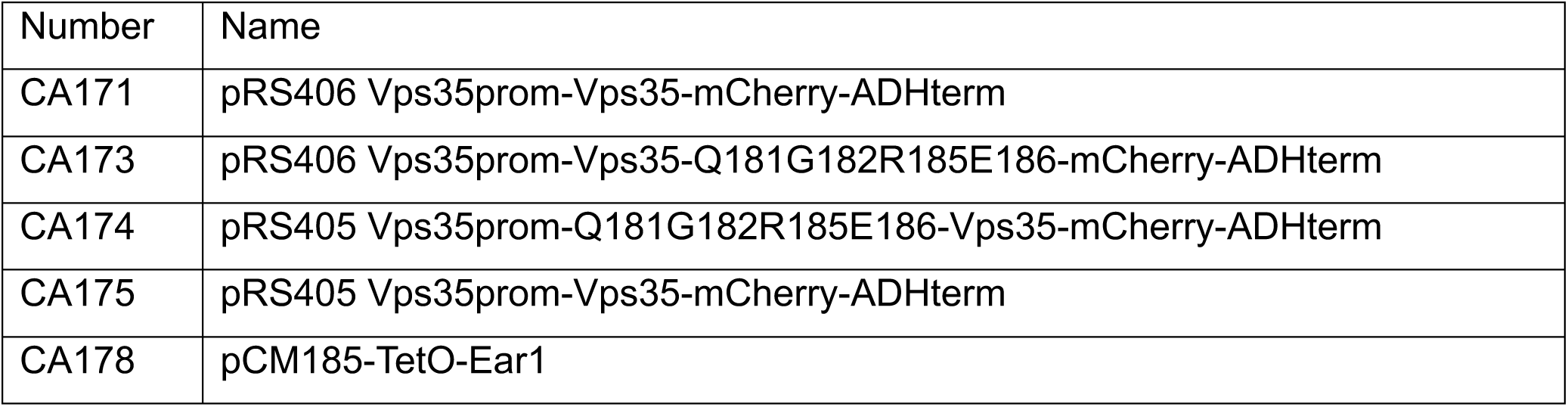
Plasmids used in this study.

**Appendix table 3:**
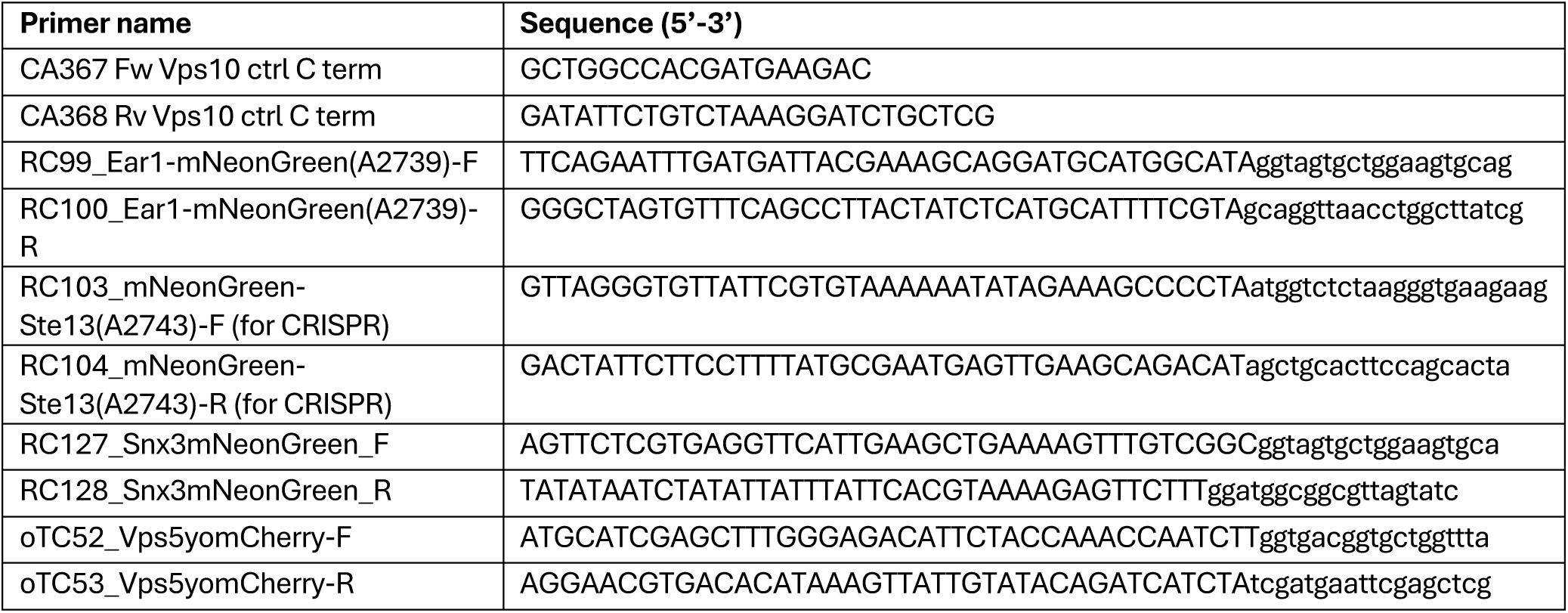
Primers used in this study.

## Acknowledgements

This work has been supported by grants from the Swiss National Science Foundation to AM (179306 and 204713), and by EMBO to SRC (ALTF 240-2023).

## Figure legends

**Supplementary Figure 1:**
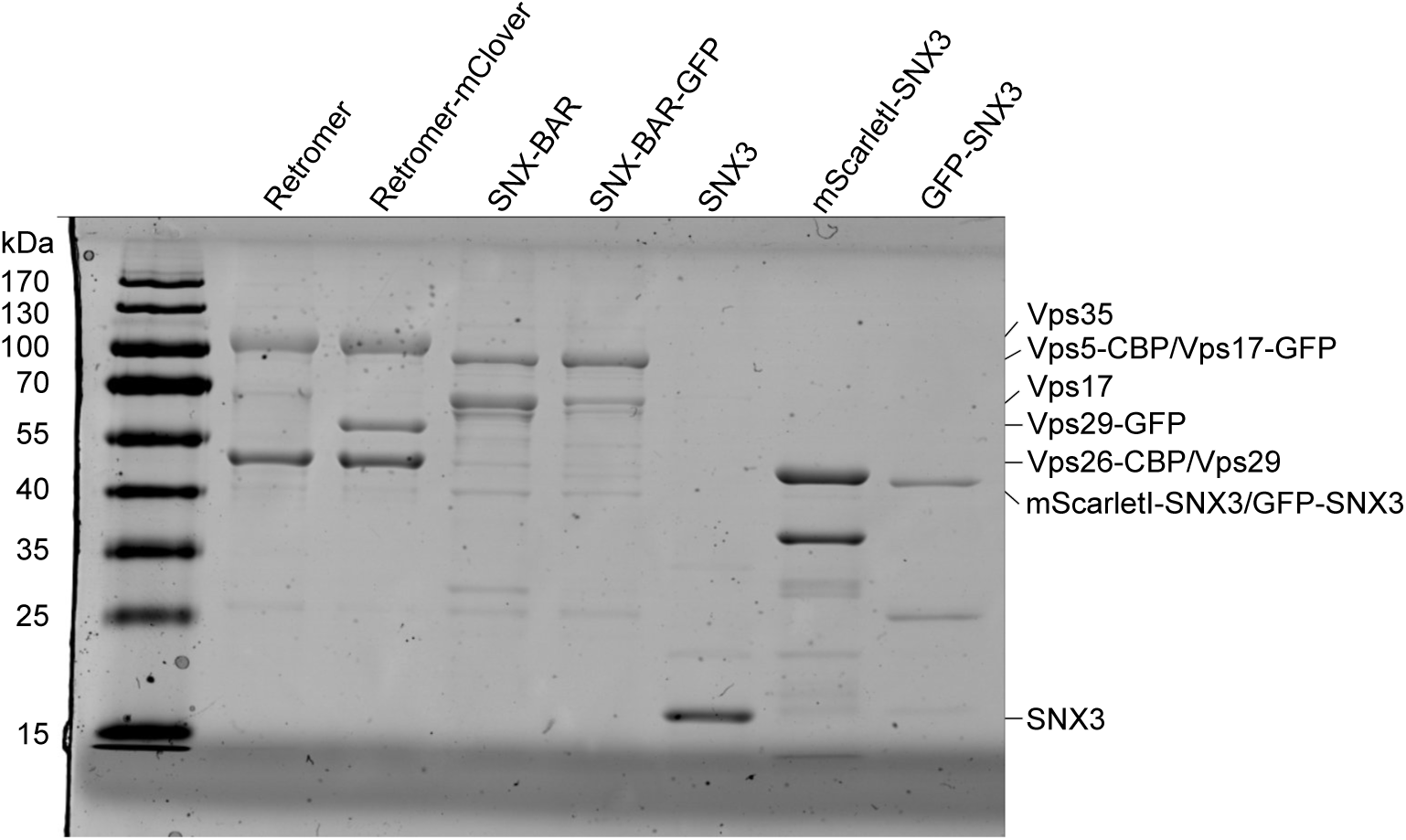
Coomassie-stained SDS-PAGE gel of the protein preparations used in the in vitro experiments in Figures 1 to 6.

**Supplementary Figure 2:**
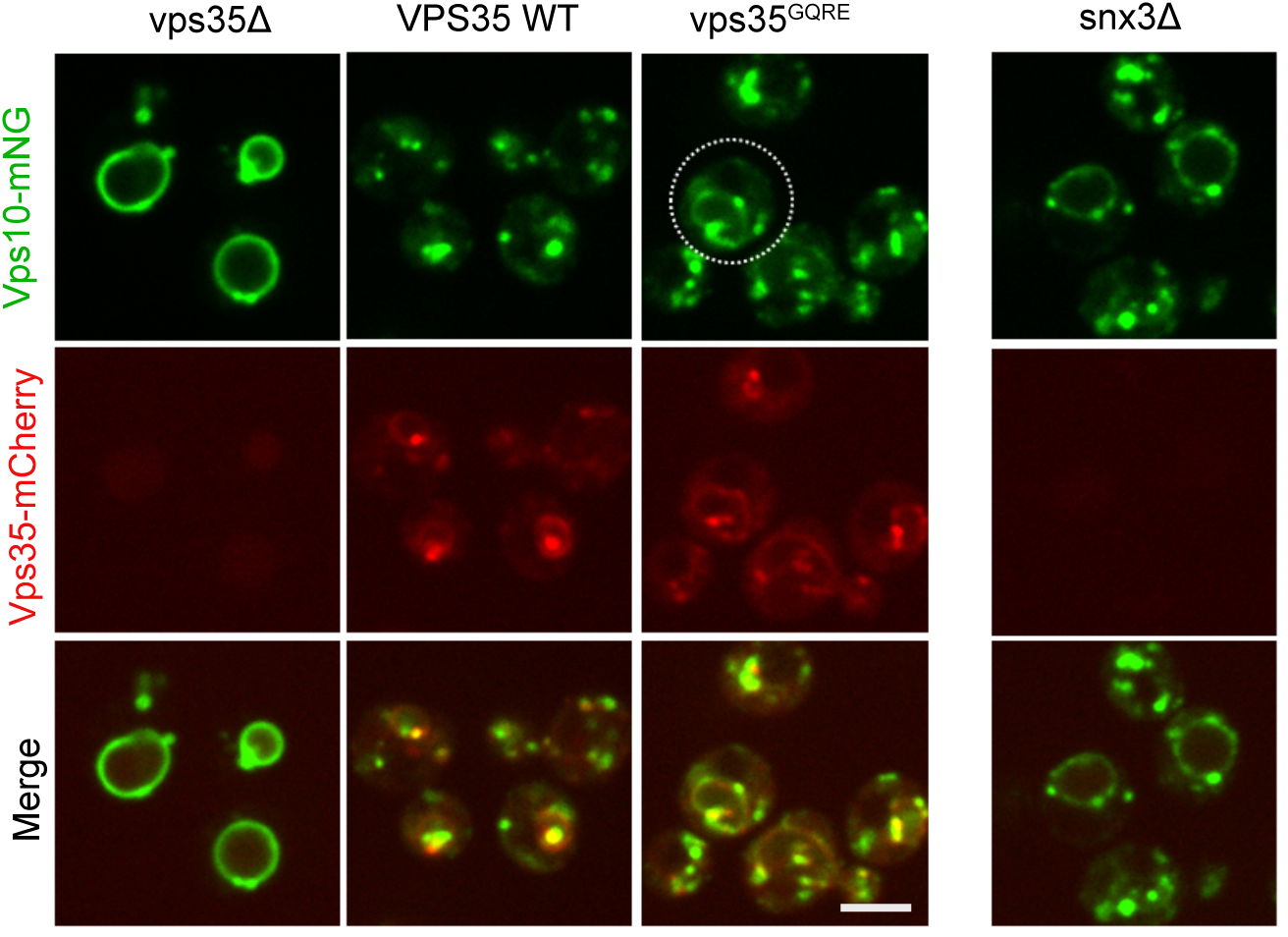
Mislocalization of Vps10 in *vps35^QGRE^* cells. Logarithmically growing *vps35Δ* cells expressing genomically tagged Vps10^mNG^ were transformed with plasmids expressing Vps35^mCherry^ (WT), Vps35^QGRE-mCherry^, or nothing. The cells were logarithmically grown overnight and analysed by spinning disc microscopy. Scale bar: 5 µm

